# Natural selection shapes and regulates the population dynamics of birds and mammals

**DOI:** 10.1101/2021.11.27.470201

**Authors:** Lars Witting

## Abstract

I fit species-specific age-structured population dynamic models to 3,369 and 483 abundance timeseries for 900 and 208 bird and mammal species, using AIC model-selection to differentiate dynamics with and without natural selection. The AIC includes natural selection in 79% to 92% of the accepted models, explaining 80% of the population dynamics variance, with median selection regulation estimated 1.2 (se:0.11) times stronger than density regulation. The estimated regulation is naturally selected by the population dynamic feedback selection of the density-frequency-dependent interactive competition among the individuals in the population, and it accelerates and decelerates population growth around the naturally selected population dynamic equilibrium, generating damped to stable population cycles with median periods around seven and eight generations given stable cycles. These results resolve several population cycle enigmas, highlighting the population dynamic implications of a feedback selection that predicts also the life history and abundance body mass allometries of animals. The selection regulated models are available for online simulations at https://mrLife.org.

## 1 Introduction

Although often neglected, a widespread absence of fit between standard population dynamic models and timeseries of abundance data remains one of the most enduring problems in population ecology (Myers 2018; Oli 2019; Andreassen et al. 2021). This reflects a paradox where population regulation is supposed to shape the dynamics of natural populations, but traditional density regulated models fail to explain even simple population dynamic trajectories. Take for example a population that declines until the decline gradually stops and the population begins to increase. While this is observed often, density regulated populations will only increase, or decline, towards carrying capacity showing no change in the direction of growth (over-compensation from strong density regulation does not explain a gradual change in the direction of growth).

For the last century or so, variation outside of the population—in the environment and in the regulating interactions with other species—provided the conceptual solution to the lack of fit between density regulation theory and data. This variation explains population dynamic deviations that correlate with interspecific interactions, environmental fluctuations, and climatic change. These drivers are case specific, and the vast majority of the available timeseries of abundance data have no identified external cause. The extend of the external influence is therefore unresolved in most cases, with the extrinsic hypothesis serving as a convenient conceptual solution where presumed ecosystem interactions disrupt the fit of single-species density regulation theory to data.

Delayed density regulated models provide the practical solution where “single-species” models “explain” much of the observed dynamics (e.g. Turchin and Taylor 1992; Hörnfeldt 1994; Hansen et al. 1999). But there is usually not an identified mechanism of cause and effect for the estimated delays, with most of the delayed density regulated studies turning the blind eye to the real problem: the widespread lack of identified inter-specific interactions that explicitly explain the observed dynamics.

If we concentrate on studies with a mechanistic focus, population dynamic explanations have advanced over the last 30 years by spatial synchrony (Ranta et al. 1995; Koenig 2002; Liebhold et al. 2004), stochasticity (Kaitala et al. 1996; McKane and Newman 2005), environmental oscillations (Post and Forchhammer 2002; Taylor et al. 2013), maternal effects (Ginzburg 1998), demographic details (Murdoch et al. 2002; McCauley et al. 2008), and higher-dimension interactions (Tyson et al. 2010; Liu et al. 2013). But even this broader theoretical framework leaves several of the most essential population dynamic enigmas unsolved (Myers 2018; Oli 2019; Andreassen et al. 2021).

Unresolved issues include the presence of population cycles in the absence of cyclic species interactions. The *Daphnia*-algae study of Murdoch and McCauley (1985) is a well-documented example, where both laboratory and natural populations of *Daphnia* cycle with a relatively constant period independently of the presence versus absence of a cycle in the algae. Similar observations have been made for snowshoe hares that cycle in the absence of lynx (Keith 1963), and there is no firm predator-prey interaction for one of best documented cycles in forest insects (Berryman 1996).

Another paradox is the widespread presence of life history changes in phase with population dynamic cycles. These changes do not follow from density regulation, and nor from interactions between predators and prey: Where predation affects survival, “most, if not all, cyclic rodent populations are characterised by phase related changes in body mass, social behaviour, … and reproductive rates” (Oli 2019). These life history cycles pose a serious problem for traditional population regulation theory because the reproductive rate stays low during the low phase of population cycles, where relaxed density regulation predicts a high reproductive rate (Myers 2018).

Other problems include that no experimental manipulation of predators and resources “has succeeded in stopping rodent population cycles anywhere” (Oli 2019), and “how can low amplitude cycles persist if high densities are required for the build-up of predators, parasitoids, pathogens or detrimental conditions” (Myers 2018).

While none of these issues question external factors as important for population dynamics, they hint at a theory that lacks an essential population dynamic mechanism. In my search for such a mechanism I take the parsimonious view that to explain the growth, abundance, and dynamics of natural populations we need first to understand how they regulate their own growth when other things are equal. Theoretically, it is not only density regulation, but also the population dynamic feedback selection of the density-frequency-dependent interactive competition in the population that regulates the growth and abundance shaping the dynamics of natural populations (Witting 1997, 2000b). Natural selection may thus— dependent upon its strength of regulation—be a main reason for the often-observed lack of fit between density regulation theory and data.

### 1.1 On natural selection regulation

With the Malthusian parameter *r* being the exponential growth rate of the population and the natural selection fitness of the individual (Fisher 1930), natural selection acts stronger on the demographic components of population dynamic growth than on other phenotypic traits. Traditional natural selection concepts, however, are typically not regulating population dynamic growth as they are population dynamically unbalanced with no feedback selected regulation. The classical example is the frequency-independent selection of traditional life history theory (Roff 1992; Stearns 1992; Charlesworth 1994) that selects a continued increase in the population dynamic growth and carrying capacity of populations (Roughgarden 1971; Caswell 1989), following Fisher’s (1930) fundamental theorem of a natural selection increase in the Malthusian parameter (Witting 2000a). This predicts a world of famine and overexploited resources (Malthus 1798) that leaves the population regulation of a balanced world outside of the domain of natural selection, in the hands of the density regulating population dynamic interactions among species (Hairston et al. 1960; Sinclair 1989; Hairston and Hairston 1993).

The counterexample is the arms race selection of a strong intra-population, density-invariant, and frequency-dependent interactive competition that selects a continued increase in competitive quality with a trade-off decline in the Malthusian parameter that drives the population to negative growth and extinction (Simpson 1953; Parker 1983; Haigh and Rose 1980; Maynard Smith 1982; Vermeij 1987). While acting in the opposite direction, this selection is just as unregulatory and population dynamically unbalanced as frequency-independent selection.

None of these selections are likely to operate on their own as natural populations seem not to be selected towards infinite growth/abundancies or towards extinction, but towards life histories that are balanced with intermediate growth and intermediate population densities. To obtain this balance we need a regulatory selection where the selection increase in population growth and carrying capacity reverts to a selection decline when the population grows beyond a certain threshold. This change in the direction of selection is indicated by arms race models as the frequency-dependent interactive competition among the individuals in the population is density-dependent, instead of density-invariant as assumed.

This density dependent interactive competition is commonly observed in natural populations across the animal kingdom (Hardy and Briffa 2013), producing a density-frequency-dependent feedback selection (Witting 1997, 2000b) where the number of competitive encounters per individual is positively related to the density of individuals in the population, and the frequency distribution of the competitive abilities/qualities across individuals makes the competitive ability/quality of an individual relative by determining the percentage of the interactive competitive encounters that the individual wins. Combined with the energetic quality-quantity trade-off (Smith and Fretwell 1974; Stearns 1992) between competitive quality and the rate of reproduction, it provides a population dynamic feedback tension where the frequency-independent selection increase in intrinsic reproduction generates a trade-off decline in competitive quality, and the resulting population dynamic growth and density-frequency-dependent interactive competition selects an increase in competitive quality with a trade-off decline in reproduction.

The population dynamic feedback between these two counteracting forces of selection regulates the population, selecting a balanced life history with limited population growth and an equilibrium abundance where the level of interactive competition is selected to be exactly so high that the resource bias of interactive competition counterbalances the selection against quality by the quality-quantity trade-off (Witting 1997, 2000b, 2002a).

This inclusion of intra-specific interactive competition alters natural selection from “a selection increase in average fitness” (Fisher 1930) to “a selection change in relative fitness” (Witting 2026), a transition that is necessary and sufficient to explain the evolutionary succession of the major lifeforms from the origin of replicating molecules (Witting 1997, 2008, 2026).

The origin selects net energy for replication, and the resulting population growth and interactive competition reallocation-selects the selected net energy into competitive traits like multicellular body mass and non-replicating interacting males in sexually reproducing units, explaining evolutionary transitions from virus-like replicators, over prokaryote- and protozoa-like self-replicating unicells, to multicellular sexually reproducing animals (Witting 2002b, 2008, 2017b, 2026).

With the intra-specific interactive competition being commonly observed (Hardy and Briffa 2013) and apparently necessary for our ability to explain the evolution of balanced animal life histories and abundancies (Witting 2008, 2026), the population dynamic question is not really whether natural selection regulation occurs, but whether it is sufficiently strong and fast to shape the dynamic of natural populations.

### 1.2 On selection regulated dynamics

Early work on the population dynamic implications of these naturally selected changes in the growth rate were verbal formulations relating primarily to the generation of population dynamic cycles (e.g. Elton 1924; Ford 1931; Voipio 1950; Chitty 1960; reviewed by Voipio 1988). These views failed to reach general consensus partially because the proposed process in the minds of many biologists, but not in the original formulations (Chitty 1996), became associated with the Wynne-Edwards (1959, 1993) hypothesis of self-regulation by group-selection, which was forcefully refuted (Maynard-Smith 1964; Wiens 1966; Williams 1966). The proponents also failed to develop population dynamic models that demonstrated the plausibility of the hypotheses (Voipio 1988), with a genetic model-interpretation with two phenotypes and no time-lag in the selection response failing to produce population dynamic cycles (Stenseth 1981, 1985).

But there is typically a delay of one generation from parents to offspring in the population response to natural selection making population dynamic cycles by the feedback selection of interactive competition theoretically plausible (Witting 1997, 2000b). These cycles depend on the magnitude of the selection response, and Sinervo et al. (2000) confirmed their existence by observing interactive competition for a two-strategy two-generation selection-driven oscillation in side-blotched lizard (*Uta stansburiana*). Other empirical studies soon documented a wide range of population dynamic implications of natural selection (e.g. Yoshida et al. 2003; Bell and Gonzalez 2009; Coulson et al. 2011; Turcotte et al. 2011; Bell 2017; Halley et al. 2021; Pavithran and Sujith 2022). Where these studies deal with specific mechanisms in a few selected species, I analyse 3,852 population dynamic timeseries covering 1, 108 species to estimate if population dynamic feedback selection is an important and widespread population regulation mechanism that will make our single species model work as a first approximation of the population dynamics of birds and mammals.

To estimate the population dynamics of natural selection changes in the growth rate, I follow a long tradition where the theoretically expected density regulated decline in *r* with increased abundance is incorporated into the structure of population dynamic models that are used to statistically estimate a parameter for the strength of the density regulation response in timeseries of abundance data (e.g. Turchin and Taylor 1992; Sibly et al. 2005; Knape and de Valpine 2012). Following Witting (1997, 2000b, 2013) I include also the theoretically expected selection regulation in the model structure allowing me to estimate an extra parameter for the strength of natural selection regulation. For this I extend the following discrete version

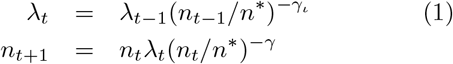

of Witting’s (1997, 2000b) selection regulated population dynamic model to species-specific age-structured models, with 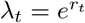 being the intrinsic per-generation replication rate in generation *t, n* the abundance with superscript * denoting population dynamic equilibrium, *γ* the density regulation parameter that defines realized replication as *λ*_*t*_(*n*_*t*_*/n*^*^)^*−γ*^, and *γ*_*ι*_ the selection regulation parameter that specifies the selection response of competitive quality (*q*) and intrinsic growth (*λ* = 1*/q*), with *λ*^*^ = *q*^*^ = 1.

One way to think of the difference between density and selection regulation is that density regulation accounts for the average suppression of the exponential growth rate *r* by the joint effects of exploitative and interactive competition, while selection regulation accounts for the average per generation selection response of the growth rate to the intra-population variation in the growth rate across the life history variants in the population.

The population dynamic equation (eqn 1) deals with the population averages of these responses, with the structural form of the selection response being theoretically deduced from the selection gradient of interactive competition across the individuals in the population (Witting 1997, 2000a,b, 2002a). If the environment is stable, most of the long-term dynamics will converge on the population dynamic equilibrium of the Continuously Stable Strategy (CSS, Eshel 1983; Taylor 1989; Christiansen 1991) selection attractor that balances the frequency-independent selection increase in *r* against the decline in *r* selected by the density-frequency-dependent interactive competition.

These interactions select for enhanced competitive quality and decelerated population growth when the population density is above the naturally selected population dynamic equilibrium, and for decreased quality and accelerated growth when the population is below the equilibrium, generating population cycles that converge on hyperexponential growth at zero abundance (Witting 1997, 2000a,b). The predicted cycles are phase-forgetting having amplitudes and cyclic regularities that depend on the magnitudes and regularities of external perturbations, with amplitude damping inversely related to the population response to feedback selection (Appendix A.2). The resulting dynamics can imitate the population cycles of forest insects (Witting 1997, 2000b), the delayed recovery of large whales following commercial exploitation in past centuries (Holt 2004; Witting 2013), and a wide range of the observed dynamics in birds and mammals (Witting 2024a).

### 1.3 Estimating regulation strengths

Several studies have estimated the strength of density regulation from statistical analyses of timeseries of abundance data (e.g. Turchin and Taylor 1992; Sibly et al. 2005; Knape and de Valpine 2012), but no study has systematically estimated the influence of selection regulation across a large number of species. I aim to estimate both regulations as it is possible to distinguish between them statistically when we fit population models to timeseries of abundance estimates. Following the theoretical expectations of eqn 1, density regulation sets the growth rate as a monotonically declining function of density, while feedback selection accelerates and decelerates growth as a function of the density frequency dependent selection (Witting 2000b). I use this mechanistic difference to provide the first large-scale analysis that estimates the strength of density and selection regulation across almost four thousand populations of birds and mammals.

My study estimates the strengths of regulation statistically from timeseries of abundance data, providing no direct observable evidence of the underlying density and selection regulating mechanisms. In addition to the uncertainty on the details of density regulation, the uncertainty includes the heritability of the selection response, which is unknown except that it is part of the complete general inclusive inheritance system of the population (Mameli 2004; Danchin et al. 2011). Population genetics is the traditional inheritance component, with other potentially important components including cultural inheritance (Danchin et al. 2011; Whitehead et al. 2019), parental effects (Boonstra and Hochachka 1997; Inchausti and Ginzburg 2009), epigenetic inheritance (Richards 2006; Bossdorf et al. 2008), and long-term selected phenotypic plasticity that allows individuals to respond to e.g. cyclic changes in the selection pressure (Snell-Rood et al. 2018; Pfenning 2021). To statistically estimate the cumulated response of these processes I assume continuous quantitative traits and a proportional additive response of the overall heritability to the selection pressure, producing a response that is similar in structure to that of the secondary theorem of natural selection (Robertson 1968; Taylor 1996; see appendix for details).

As it is the cumulated response of the inclusive inheritance system to the selection pressure that generates the change in population dynamic growth, we cannot reject selection regulation as a valid hypothesis should one inheritance component be insufficient to account for the observed change. Hence, I treat the estimated models as we usually treat statistical estimates of density regulation. From our theoretical reasoning we expect both density and selection regulation in natural populations knowing how they affect the dynamics structurally, and the implications of the statistical estimates are therefore accepted as our best model estimates of the dynamics, acknowledging that the underlying mechanistic details are not directly observed.

### 1.4 Species specific models

To get as accurate estimates of density and selection regulation as possible I use species-specific age-structured models that I obtain from an inter-specific interpolation independently of the timeseries of abundance estimates. For this I use the theoretically predicted inter-specific body mass variation and life history allometries (Witting 1995, 2017a) of the feedback selected population dynamic equilibrium of eqn 1 in an inter-specific allometric trait interpolation across more than hundred thousand life history data, estimating age-structured equilibrium population dynamic life history models for approximately 90% of the birds and mammals of the world (Witting 2024a).

I use these species-specific life history models as the equilibrium age-structured population dynamic models of the present study. These models project a stable population with zero growth, and by including the population regulation mechanisms of eqn 1 I can focus my timeseries analysis on the estimation of the population regulation parameters and initial conditions that explain the observed dynamics.

## 2 Method

### 2.1 Data

I use timeseries of abundance estimates from the Living Planet Index (LPI 2022), the North American Breeding Bird Survey (BBS; Sauer et al. 2017), the PanEuropean Common Bird Monitoring Scheme (EU; PECBMS 2022), the Netwerk Ecologische Monitoring (NET; Sovon 2022), the Swiss Breeding Bird Index (SWI; Knaus et al. 2022), the British Trust for Ornithology (BTO 2022), the Danish Ornithological Society (DOF 2022), and Svensk Fågeltaxering (SWE; SFT 2022).

Having different origins, these timeseries are of varying quality and length. To analyse a large number of timeseries I apply a moderate baseline filter to the data, and to identify results from long timeseries of high-quality data I select a restricted subset of controls. The baseline includes timeseries with more than ten estimates over at least fifteen years, resulting in 3,369 accepted timeseries for 900 bird species and 483 timeseries for 208 mammal species, with all timeseries scaled for a geometric mean of unity.

The high-quality controls are a subset of the available standardised indices from point-counts of birds. These indices are calculated from hundreds of separate indices for individual observers that count in the same way at the same time each year on individual routes with a given number of geographically fixed point-counts. The calculation of these indices is very standardised, correcting for observer and severe weather, providing high quality timeseries given enough routes.

A potential issue with bird indices is that their geographical coverage may not necessarily reflect a population with spatially synchronised dynamics. I account for this by restricting my control timeseries to the population dynamics delineated indices (PDDIs) that Witting (2023b) compiled from the North American Breeding Bird Survey (Sauer et al. 2017). These are based on more than six million observations from USA and Canada, providing yearly abundance estimates for 51 years. Starting from a 15*x*7 longitudinal/latitudinal grid of separate indices covering USA and southern Canada, the PDDIs combine neighbouring areas with synchronised indices delineating larger areas with desynchronised dynamics from one another, estimating 462 populations with different dynamics (Witting 2023b).

### 2.2 Population models

I use species-specific age-structured models to ensure that the estimated regulation and dynamics occur on the relevant biological timescale accounting for the demographic delays and population dynamic inertia of each species. If I used a non-structured replication model (like eqn 1) with yearly timesteps instead, I would convert much of the between year fluctuations in the abundance estimates into strong density regulation in over-compensatory models (Wolda and Dennis 1993). This may happen even when the true population trajectory is smooth, and the fluctuations in the abundance estimates follow from the estimation process only. Such fluctuating sampling variation is widespread as it is common to count only fractions of natural populations with year-to-year variation in the available fraction.

To reduce overfitting to random sampling variation further, I focus on smooth population trajectories by restricting the estimated strengths of density and selection regulation to weak and moderate levels to avoid false estimates of over-compensation (see appendix), allowing additional sampling variation to account for potential year-to-year zig-zag variation in the data (see appendix). This maintains the estimated regulation within realistic limits, assuming that truly fluctuating dynamics is rare at the level of bird and mammal populations.

As we cannot usually estimate the age-structured life history from abundance data, I obtain the equilibrium demographic parameters of the age-structured models of all species from Witting (2024a). These parameters are offspring survival [*p*_0_ for age-class (*a*) zero], annual survival (*p*) for older age-classes (*a* ≥ 1), age of reproductive maturity 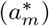, and annual reproduction (*m*^*^) of mature 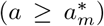 females at population dynamic equilibrium (superscript *).

These models have a species-specific equilibrium age-structure (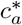 with 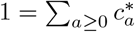) that I use as the initial age-distribution, providing a complete equilibrium age-structured population dynamic model with zero growth that I can iterate forwardly in age-unit-timesteps (often years but see appendix). The equilibrium per-generation replication rate

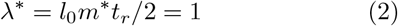

is a condensed expression of each model where *r*^*^ = ln *λ*^*^ = 0, with *t*_*r*_ = 1*/*(1 *− p*) being the expected reproductive period of offspring that survive with probability 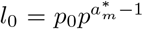 to reproductive maturity (Witting 2024a). By keeping these equilibrium demographic parameters fixed for each species, I estimate only the absolute abundance, the density and selection regulation parameters, and a few initial conditions from the abundance data.

I develop exponential, hyperexponential, density regulated, and selection regulated models for each population (see below), using model selection to obtain an initial estimate of the dynamics. For this I use likelihood fitting based on log normally distributed abundance data and apply a minimum fitting criterion (Appendix A.3) to find the best-fitting-hypothesis by the Akaike information criterion (AIC, Akaike 1973) to trade-off the number of parameters (from 2 to 5) against the likelihood of each model (Appendix A.3).

As this AIC selects selection-based models most often (see results), I run a second AIC-selection to estimate the best selection regulated models for all populations. In addition to the stable equilibrium of the initial model, this second selection includes also models with a linear trend in equilibrium density during the whole, or parts of, the data period (Appendix A.3). This allows me to quantify not only the relative strengths of regulation by density and selection, but to estimate also if local population trends are indicators of underlying changes in the external environment (assuming that a change in equilibrium reflects improved or deteriorating external factors).

I set regulation by density and selection to operate on the birth rate 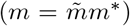 and age of reproductive maturity 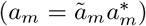 by changes in relative parame-ters (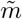 and 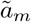). These are constant in the exponential population model and unity (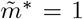 and 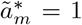) at the population dynamic equilibrium (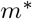 and 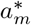) of the regulated models.

As I fit the 1+ abundance (*n* = Σ _*a*≥1_ *n*_*a*_) of the population to the abundance data, the estimated multiplicative regulation on the birth rate includes regulation on offspring survival implicitly (as the number of age-class one individuals at time *t* is the number of births at time *t* − 1 multiplied by offspring survival). Hence, I cover regulation on the three life history parameters that are most sensitive to density dependent changes, allowing for regulation on *a*_*m*_ for an extended analysis of the PDDI control timeseries only. Appendix A.1 describes the details of the age-structured population dynamic modelling based on selection regulation, with essential differences between the four population models described below.

#### Exponential growth

The exponential models are the life history models with *p*_0_, *p*, 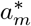, and *m*^*^ obtained from Witting (2024a), *ã*_*m*_ = 1, and relative birth 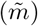 and initial abundance (*n*_*t*=0_) at time-step zero (*t* = 0) estimated from data.

#### Hyperexponential growth

The age-structured abundance 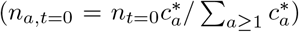 at time zero is the only initial condition of the exponential and density regulated models. A uniform *q*_*a,t*=0_ = *q*_*t*=0_ vector of competitive quality (*q*) by age is an additional initial condition in the selection models where the projection of offspring quality (*q*_0,*t*_) is given by selection acting on the quality of parent (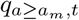; Witting 1997, 2000b).

I use the competitive quality parameter *q* as a joint measure of the energy requiring traits that intra-specific interactive competition selects to enhance the competitive ability of the replicating unit (traits like body mass, kin group size of cooperating sexually reproducing individuals, and interactive behaviour; Witting 1997, 2017b). The energetic quality-quantity trade-off

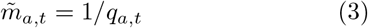

is then defining the relative birth rate, with time used to build quality defining relative reproductive maturity (ã_*m*_) proportional to quality

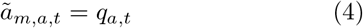

when included as a selection response in the extended analysis of the PDDI data, with *q*^*^ = 1 for all *a* at equilibrium with no growth.

The population dynamic selection response of my model follows from a structural analysis of selection across the intra-population phenotypic variation (subscript *i*) in individual competitive quality covering the intra- and inter-age-class variation implicitly. Following Witting (1997, 2000b), I calculate the selection gradient on log scale (*∂r*_*i,t*_*/∂* ln 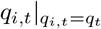) at the average (*q*_*i*_ = *q*) quality of the population. Then, by extending the logic of the secondary theorem of natural selection (Robertson 1968; Taylor 1996) to inclusive inheritance (Mameli 2004; Danchin et al. 2011), the selected change in ln competitive quality—and thus indirectly in reproduction and reproductive maturity by eqns 3 and 4—is approximated as a product

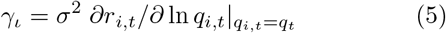

between the selection gradient and the additive heritable variance (*σ*^2^ ≥ 0) of the inclusive inheritance system (Appendix A.1), with *γ*_*ι*_ being a time invariant selection response for the hyper-exponential model, with the imposed limits on *γ*_*ι*_ examining weak to moderate selection responses only.

When there is no interactive competition and all individuals have equal access to resources, the intra-population variation in the growth rate is *r*_*i,t*_ ∝ − ln *q*_*i,t*_ from eqns 1, 2, and 3, with a time invariant selection gradient of *∂r*_*i,t*_*/∂* ln 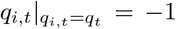 with *γ*_*ι*_ = −*σ*^2^. This is the limit case of hyperexponential growth at zero population density, selecting for a decline in *q* and increase in *r*. Yet, I allow for both positive and negative *γ*_*ι*_ values to capture constantly accelerating (*γ*_*ι*_ *<* 0) and decelerating (*γ*_*ι*_ *>* 0) growth (*γ*_*ι*_ = 0 is exponential growth). As the selection gradient on the per-generation growth rate is 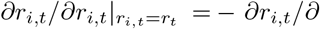 ln 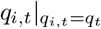, the acceleration/deceleration of the growth rate is

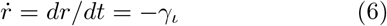

With the selection response acting on log scale, the average offspring quality (*q*_0_) at time *t* is a product

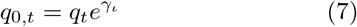

between the time invariant selection response 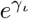 and the average quality of the mature individuals

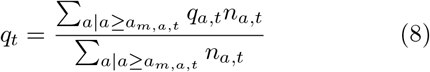

With eqns 7 and 8 capturing the response of selection for the age-structured population model, it is not necessary to include the intra- and inter-age-class variation in competitive quality and fecundity explicitly into the calculation of the selection response. Hence, I avoid dealing with complex individual-based models to track the differentiation in competitive qualities, resource availabilities, and reproductive rates across all the individuals in the population.

The hyperexponential model is structurally more complex than the exponential, yet it has only a single parameter (*γ*_*ι*_) and two initial conditions (*n*_*t*=0_ & *q*_*t*=0_) to estimate from data.

#### Density regulated growth

The density regulated models resemble the exponential models, but they have a Pella and Tomlinson (1969) regulated relative reproduction

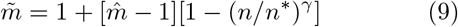

with three parameters (maximum relative birth rate 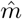; density regulation *γ*; equilibrium abundance *n*^*^) and one initial condition (*n*_*t*=0_) to estimate from data.

#### Selection regulated dynamics

The selection regulated models are structurally similar to the hyper-exponential models, but they have density and selection regulation acting on relative reproduction (and also on relative age of maturity for the extended PDDI analysis). This density regulation is a log-linear perturbation

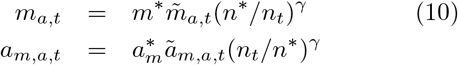

of the life history that evolves (changes in 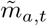 & *ã*_*m,a,t*_ by eqns 3, 4, and 11) around the evolutionary equiLibrium 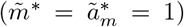. Having a life history that is selected/adapted to the resource level at population dynamic equilibrium, the density regulation function of the selection regulated model operates as a local approximation around the equilibrium density, lacking the maximum growth (*r*_*max*_) / birth 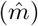 rate parameter that is defined explicitly for zero density in traditional density regulation models (eqn 9).

Following Witting (1997, 2000b), Appendix A.1 derives the change in competitive quality by the population dynamic feedback selection of density dependent interactive competition as

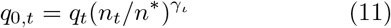

with *q*_*t*_ from eqn 8 and the selection induced acceleration/deceleration of the growth rate

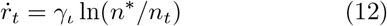

being a log-linear function of the density dependent ecology. This model has three parameters (*γ, γ*_*ι*_, & *n*^*^) and two initial conditions (*n*_*t*=0_ & *q*_*t*=0_) to estimate from data (and 1 to 3 extra parameters when *n*^*^ can change, see Appendix A.3).

Eqn 11 is based on the counteracting selection of the quality-quantity trade-off and the biased resource access from density-frequency-dependent interactive competition (Witting 1997, 2000b). The explicit modelling of the selection requires equations that account for the intra-population variation in competitive quality, the density dependence in the level of interactive competition, and the biased resource access from this competition. The population dynamic response of this selection converges on eqn 11 that describes the population level response in the population dynamic model (Appendix A.2).

The dynamics of population dynamic feedback selection is cyclic, and I calculate the cycle period (*T*, in generations) and damping ratio (*ζ*) to describe it. The damping ratio is one for the monotonic return of density regulated growth (*γ*_*ι*_ = 0 & *γ >* 0), it is between zero and one for damped cyclic dynamics (0 *< γ*_*ι*_*/γ <*≈ 1), zero for stable cycles (*γ*_*ι*_*/γ* ≈ 1), and it declines to minus one for increasingly unstable cycles with amplitudes that increase over time instead of dampening out (*γ*_*ι*_*/γ >*≈ 1, see appendix).

## 3 Results

Population models for 2,058 bird and 290 mammal populations passed the minimum fitting criterion during the first round of AIC model selection. This model selection selected selection-based models in 79% of the cases for both birds and mammals, with selection regulated models selected in 43% of the cases, followed by 35% hyperexponential, 14% exponential, and 7.6% density regulated models.

The selection effects were more pronounced in the PDDI control timeseries. Here, AIC included selection in 92% of 267 chosen models, with selection regulated models selected in 69% of the cases, followed by 23% hyperexponential models, 4.9% exponential, and 3.4% density regulated models.

With selection regulation covering all models (exponential when *γ* = *γ*_*ι*_ = 0; hyperexponential when *γ* = 0 & *γ*_*ι*_ ≠ 0; density regulated when *γ >* 0 & *γ*_*ι*_ = 0), I use the second AIC selection of selection regulated models with and without a change in equilibrium to describe the dynamics. This resulted in 2,801 accepted models (2,480 for birds; 321 for mammals) that explained 80% of the variance in the data, on average.

Fig. 1 shows the estimated dynamics of 24 populations, with the Supporting Information showing all models. Where density regulated growth returns monotonically to carrying capacity (like top left plot in Fig. 1), the majority of the estimated selection regulated dynamics have damped to stable population cycles. Few populations show exploding cycles with negative damping ratios, but these may reflect estimation uncertainty.

**Figure 1:**
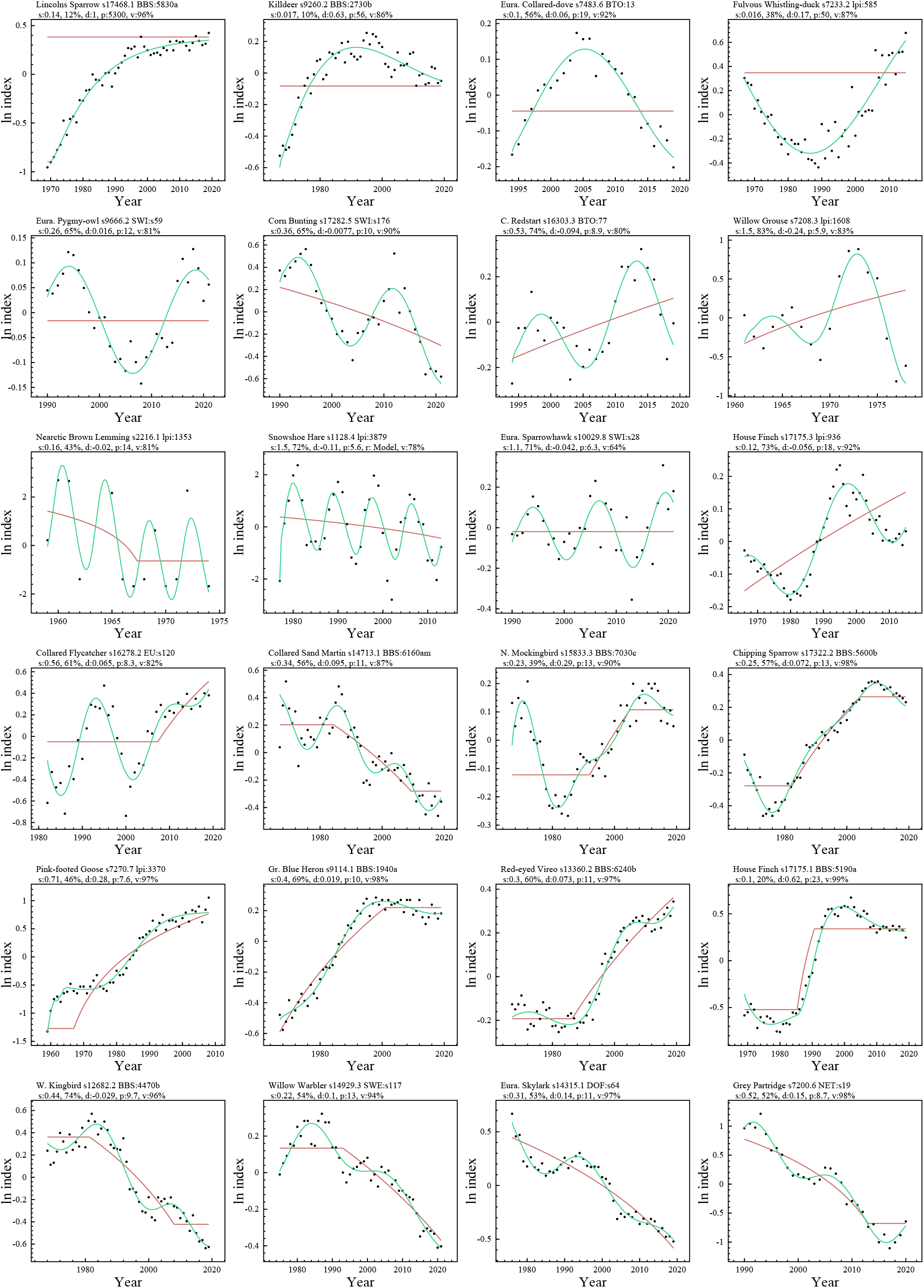
Examples of selection-regulated models fitted to population dynamic timeseries. Dots are index series of abundance, red lines the estimated equilibria, and green curves the model projections. Headings: name; id. nr; data reference; s:*γ*_*ι*_ *& γ* _*ι*_*/*(*γ* _*ι*_ + *γ*) in %; d:damping ratio; p:period in generations; v:explained variance in %.

The estimated median selection regulation (*γ*_*ι*_) is 0.36 (se:0.01) for birds and 0.72 (se:0.032) for mammals, with median density regulation (*γ*) being 0.31 (se:0.0088) for birds and 0.31 (se:0.028) for mammals. The left plots in Fig. 2 show the distributions of the relative strength of selection regulation [i.e., *γ*_*ι*_*/*(*γ* + *γ*_*ι*_)] across all timeseries with accepted selection regulated models. With median estimates of 0.55 (se:0.0058) for birds and 0.58 (se:0.017) for mammals, selection regulation is equally or more important than density regulation in most populations, with median regulation ratios (*γ*_*ι*_*/γ*) of 1.2 (se:0.11) and 1.4 (se:0.37). These results resemble those of the PDDI controls, where relative selection regulation [*γ*_*ι*_*/*(*γ* + *γ*_*ι*_)] is 0.53 (se:0.012) at the median across 399 selection regulated models. Allowing for regulation on reproductive maturity among the PDDI controls, 47% of 408 AIC-selected models were regulated through the reproductive rate and age of maturity, with a median relative [*γ*_*ι*_*/*(*γ* +*γ*_*ι*_)] selection regulation of 0.61 (se:0.019) across all models.

**Figure 2:**
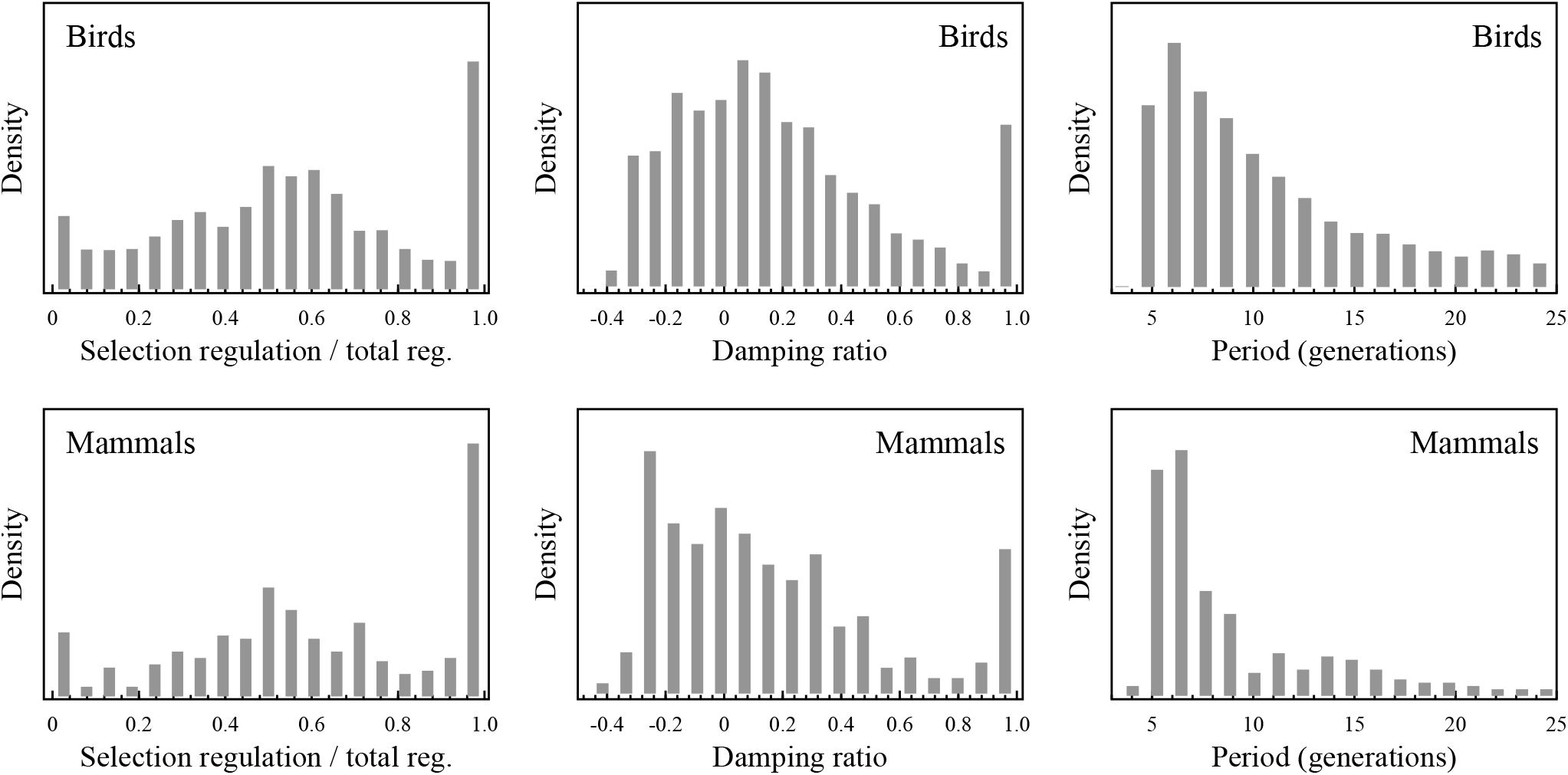
Distributions of the strength of selection regulation [divided by total regulation; *γ* _*ι*_*/*(*γ* _*ι*_ + *γ*)], the damping ratio, and population period of the final AIC selected models for 2,480 bird and 321 mammal populations.

The distributions of regulation estimates cover the range from almost pure selection regulation to almost pure density regulation. Only 7.3% of the bird and 6.5% of the mammal populations have selection regulation below 10% of total regulation by density and selection. These results cannot support the hypothesis that natural populations of birds and mammals are density regulated predominantly. With the results of the first and second round of AIC selections agreeing on limited support for pure density regulation, the structural difference (eqn 9 vs. eqn 10) in the density regulation functions of the density regulated and selection regulated models is not influencing the conclusion.

The middle plots in Fig. 2 show the distributions of the estimated damping ratios. With median damp-ing ratios around 0.12 (se:0.0071) and 0.062 (se:0.021) the population dynamics of birds and mammals is best characterised as cyclic. 83% of the bird populations, and 85% of the mammals, have damping ratios that are smaller than 0.5. Only 7.3% of the bird populations, and 7.8% of mammals, have strongly damped density regulation like growth with damping ratios above 0.9.

The right plots in Fig. 2 show the distributions of the periods of the population cycles. Only 3% of the estimated periods are shorter than five generations. The distributions have long tails toward long population periods, and they peak in the lower range with 46% of all birds, and 62% of all mammals, having periods below 10 generation. Median estimates are 11 (se:86) generations for birds and 7.9 (se:180) for mammals, and the period increases with stronger damping. The median period increases from 8.3 (se:0.99) and 7.2 (se:0.85) generations for birds and mammals with stable cycles (damping ratios around zero), to 28 (se:9.5) and 15 (se:1.8) for damping ratios around 0.8. These estimates reflect a weak to moderate non-fluctuating selection regulation that operates over several generations to complete a selection cycle.

Across the AIC-chosen selection regulated models, the estimated equilibrium abundancies increase for 30% and 21% of the bird and mammal timeseries and decline for 29% and 14%. Yet, these apparent changes in the external environment do not determine the estimated local direction of growth. Where density regulated populations tend to decline only when the equilibrium abundance declines due to a deteriorating environment, the selection regulated populations often decline about 50% of the time when the equilibrium is stable, declining, or increasing (Fig. 1, second line). For cases where the estimated trajectories are either declining or increasing locally, 76% of the population dynamic declines were not associated with an estimated decline in the equilibrium abundance, and 77% of the population dynamic increases were not associated with an equilibrium increase. In fact, 23% of the local population declines had increasing equilibria, and 27% of the local increases had declining equilibria.

## 4 Discussion

By focussing on internal population regulation and trends in the equilibrium abundance, I accounted for 80% of the variance in timeseries trajectories for 2,480 populations of birds and 321 populations of mammals.

I used the life history estimates of Witting (2024a) to construct a complete equilibrium age-structured population dynamic model for each species, with the population dynamic effects of density and selection regulation requiring the statistical estimation of only three parameters (density regulation, selection regulation, and equilibrium abundance) and two initial conditions (abundance and quality).

I used additional variance to account for random environmental perturbations and fluctuating abundance estimation, and I used trends in the equilibrium abundance to capture directional changes in external factors like habitats, resources, and predators. My models do not account for secondary phase related effects from e.g. cyclic changes in predation mortality, and I reduced their potential confounding effects by excluding models with non-random residuals from my analysis.

I used AIC-selection to choose the best population model for each timeseries, with selection-based models selected for 79% to 92% of the analysed timeseries, with median selection regulation estimated 1.2 (se:0.11) times stronger than density regulation. This identifies selection regulation as important and pure density regulation (with no selection) as the exception rather than the rule.

With selection regulation included we obtain a flexible single-species model that describes a broad range of the observed population dynamic trajectories (Fig. 1). These are generally cyclic with median damping ratios around 0.12 (se:0.0071) and 0.062 (se:0.021) for birds and mammals respectively, and population dynamic periods that increase with increased damping, with medians around 8.3 (se:0.99) and 7.2 (se:0.85) generations for stable cycles with damping ratios around zero. You can make your own species and population specific online simulations of the estimated selection regulated dynamics at https://mrLife.org.

### 4.1 Evidence of mechanisms

Apart from the similarity between the model projections and the empirical abundance trajectories, my analysis provides by itself no direct evidence of the estimated density and selection regulation. Yet, empirical studies have already documented similar selection driven population dynamics across a wide range of species. The study by Sinervo et al. (2000) documents a natural population with a density-frequency-dependent feedback selection of interactive competition, including a selection for increased competitive quality and decreased growth above the average abundance, and selection for the opposite below. Where their clear-cut two-strategy system produced a stable two-generation oscillation, the gradual selection of the continuous traits in my study typically produces damped cycles with periods that last more than seven generations.

Turcotte et al. (2011) studied the alternative case with growing populations at a low abundance and documented—as predicted by natural selection—a selection acceleration of the population dynamic growth rate by up to 40% over few generations, contrasting to exponential and density regulated growth that assume a constant or declining growth rate for similar situations. And the selection of faster-spreading Covid-19 variants (Halley et al. 2021; Pavithran and Sujith 2022) documents the limit case with a constantly accelerating growth rate and a hyper-exponential (instead of exponential) increase in abundance (Witting 2000a), as predicted for *r*-selected replicators like virus and prokaryotes (Witting 2017b).

Other interesting evidence relates to the selection accelerated growth that turns the population decline into an increase during the low phases of population cycles. This repeated change is in essence repeated evolutionary rescue, where the predicted selection accelerates the growth rate of declining populations preventing them from going extinct by turning the decline into an increase; as documented by several independent studies on evolutionary rescue (e.g. Gomulkiewicz and Holt 1995; Agashe 2009; Bell and Gonzalez 2009; Ramsayer et al. 2013; Bell 2017). Other eco-evolutionary studies document a wide array of population dynamic responses to natural selection (e.g. Thompson 1998; Yoshida et al. 2003; Hairston et al. 2005; Saccheri and Hanski 2006; Coulson et al. 2011; Hendry 2017; Brunner et al. 2019).

Another important line of evidence for selection regulated population dynamics comes from otherwise unresolved issues with population dynamic cycles. Here selection regulation predicts that the proximate cause of the cyclic dynamics is a special type of phase related life history change that does not follow from density dependence or predator-prey interactions. This prediction includes selection for lower reproduction, larger body masses, increased competitive behaviour (like aggression), larger cooperating kin groups, and more interacting males at high population densities, and selection for the opposite at low densities (Witting 1997, 2000b).

These predictions align with plenty of empirical evidence that includes a body mass cycle in the *Daphnia* experiments of Murdoch and McCauley (1985), with larger individuals occurring mainly in the late peak phase of the cycle, and smaller individuals mainly in the early increasing phase (Witting 2000b). Voles and lemmings with cyclic dynamics have similar changes in body mass (e.g. Boonstra and Krebs 1979; Lidicker and Ostfeld 1991; Norrdahl and Korpimäki 2002), and such changes are observed also in snowshoe hare (Hodges et al. 1999) and cyclic forest insects (Myers 1990; Simchuk et al. 1999).

As predicted, the percentage of males increases in small rodents when densities are high, while females predominate during the low phase (Naumov et al. 1969). Other cases of an increased male fraction with increased density include white-tailed deer (*Odocoileus virginianus*) (McCullough 1979) and northern elephant seal (*Mirounga angustirostris*) (Le Boeuf and Briggs 1977). Individuals of voles and red grouse (*Lagopus lagopus scotica*) are, again as predicted, more aggressive at high than low population densities, with red grouse having larger kin groups evolving during the increasing phase of a cycle (e.g. Boonstra and Krebs 1979; Piertney et al. 2008).

While most of these empirical studies failed to identify the underlying cause of the observed phase related life history changes, the view that they are unrelated to natural selection seems untenable as the density-frequency-dependent feedback selection of interactive competition explicitly predicts them, while density regulation and trophic interactions do not explain them (at least not straightforwardly).

As there can be no cycle in the abundance of a closed population unless there is a cycle in at least one life history parameter, the phase related life history cycles are not just secondary side effects with no relation to the underlying cause of the cycles. Being ultimately predicted by population dynamic feedback selection, they are the necessary proximate cause that drives the population dynamics. Following a perturbation of the population dynamic equilibrium, the predicted feedback cycle starts when the associated change in the level of interactive competition selects a life history change that initiates the first population cycle, with the resulting cyclic change in abundance repeating the cyclic density dependent selection that keeps the cyclic dynamics for its extended, but often damped, existence.

Low amplitude cycles are not a problem because the cyclic dynamics of selection regulation do not depend on high-density amplitudes for the build-up of predators, pathogens, or other detrimental factors. Selection regulation accelerates and decelerates population dynamic growth in smaller or larger steps around the equilibrium abundance, with strong deceleration around the peak abundance of high-amplitude cycles selecting a reproductive rate that stays low during the low phase of the cycle until the following phase of increase.

### 4.2 Implications

The estimated dynamics, and widely observed phase related life history changes, support population dynamic feedback selection by interactive competition as a common regulator of population dynamics, in addition to density regulation. This has several population dynamic implications that make sense in a broader eco-evolutionary perspective.

A partial decoupling of population growth from the state of the environment is one intriguing consequence of selection regulation. Pure density regulation with no selection implies that a gradual change from a growing to a declining population, or from a declining to a growing population, requires a corresponding change in the extrinsic environment. This have led to the concept of indicators, where the population dynamic behaviour of indicator species is supposed to reflect the underlying state of the environment (e.g. LPI 2022; PECBMS 2022). While environmental changes may change the direction of growth, the direction may also change by selection in the absence of environmental change. In other words, the growth of populations are not necessarily indicators of the environment. For about 75% of the estimated local population declines in this study I estimated no deteriorating environment.

Other essential results relate to the population dynamic equilibrium, where the population dynamic feedback of interactive competition selects the population dynamic equilibrium abundance as an integral part of the selection attractor that determines the life history of a species (Witting 1997, 2008, 2026). This selection scales with the selected net energy of the organism, and it explains part of the observed inter-specific variation in abundance as an allometric response to the naturally selected variation in net energy and mass (Witting 1995, 2023a), resolving issues with a density regulation theory that seems not to explain the abundance of animals (May 2020).

The life history and abundance of the naturally selected population dynamic equilibrium is fundamentally changing our understanding of population regulation. This selection attractor selects net assimilated energy into population growth until the population dynamic equilibrium has evolved to an abundance that generates a level of intra-population interactive competition where the monopolisation of resources by the larger-than-average individuals counter-balances the quality-quantity trade-off that selects for smaller masses. This naturally selected distribution of resources across the individuals in the population makes the different body mass variants approximately equally fit, adapting the life history to the resource level at the naturally selected population dynamic equilibrium (Witting 1997).

The selection regulation of a naturally selected population dynamic equilibrium implies a selection that rescales density regulation by selecting the life history and physiology that matches zero population growth (*r*^*^ = 0) at the naturally selected population dynamic equilibrium. With zero population growth being the naturally selected life history optimum, selection is actively removing the famine of the traditional density regulation concept by selecting the life history and physiology that matches the available resources. This selects density regulation as primarily a local density response in the surroundings of the selected equilibrium, breaking down the assumed *r*_*max*_ to *r*^*^ tension of the traditional density regulation concept, which requires a rarely to never observed famine to supress optimal growth (*r*_*max*_) at zero abundance to zero growth (*r*^*^) at equilibrium. The selection also removes the power of regulating the long-term equilibrium abundance from the density dependent interactions among species, selecting a balanced nature where the world is green, resources are common, and most individuals are well fed (Witting 1997, 2008).

Population dynamics is then a temporal disturbance of the naturally selected population dynamic life history equilibrium. Here it is mainly the response of the feedback selection on the age-structured demography that determines the period of the cyclic dynamics, the magnitude of the environmental perturbation that determines the amplitude, and the strength of density regulation relative to selection regulation that determines the damping of the cycle. The population dynamic feedback of interactive competition may in this way select the life histories and abundancies of natural populations, with the eco-evolutionary interplay between the feedback selection, density regulation, and varying environment shaping the dynamics.

## Acknowledgements

I thank all who collect and publish population dynamic data, and reviewers for comments that improved my paper.

## Supporting Information

### si-plot

Population plots

### si-cpp

Population model in c++

## A Appendix

### A.1 Selection regulated model

I use the species-specific age-structured demography estimates of Witting (2024a) as the population dynamic equilibrium models of the species in my study. The equilibrium (denoted by superscript *) per-generation replication rate of unity

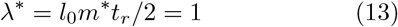

provides these models in a condensed formulation, where *r*^*^ = ln *λ*^*^ = 0 and the survival of offspring to the age of reproductive maturity 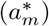 is 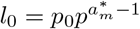, with *p*_0_ being the survival of age-class zero individuals, *p* the individual survival for older age-classes, *m*^*^ the reproductive rate of mature females (assuming an even sex ratio), and *t*_*r*_ = 1*/*(1− *p*) the expected reproductive period of individuals that mature.

The age-structured demography of Witting (2024a) is parameterised by annual estimates (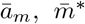, and 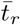 for models iterated in annual timesteps Δ*t* = 1). For species with a small ā_*m*_ and/or short 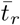 I use shorter timesteps to ensure 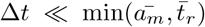, with the parameters of my age-structured iteration models being *a*_*m*_ = ā_*m*_*/*Δ*t*, 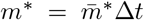, and 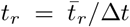, with the survival of individuals in 1+ age-classes be-ing *p* = (*t*_*r*_ − 1)*/t*_*r*_, and age-class zero survival being 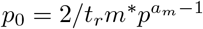 from eqn 13.

I iterate the population dynamic models at the level of the average parameter values of each age-class (subscript *a*), with subscript *a* = *x* ≫ *a*_*m*_ denoting the maximum lumped age-class. The number *n*_*a,t*_ of individuals in age-class 0 *< a < x* at timestep *t* is then

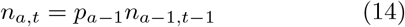

with the number in age-class *x* being

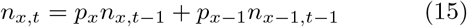

with *p*_*a*_ = *p*_0_ for *a* = 0 and *p*_*a*_ = *p* for *a* ≥ 1.

Let selection operate on the competitive qualities (*q*_*i*_) of individuals (subscript *i*), with quality defining a relative intrinsic birth rate 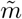 by the quality-quantity trade-off

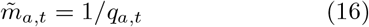

and time required to build quality defining a relative intrinsic age of reproductive maturity

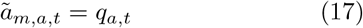

in proportion to quality, with 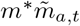 and 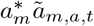 being the absolute intrinsic parameters with all individuals having 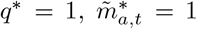, and 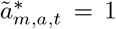 for the naturally selected population dynamic equilibrium models obtained from Witting (2024a). Assuming no change in the quality of a cohort over time, we have *q*_*a,t*_ = *q*_*a−*1,*t−*1_ and

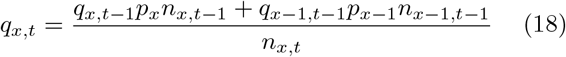

with offspring quality given by the population level selection response of eqn 27.

To formulate density dependence let the number of individuals in each age-class relate to time just after each timestep transition, with offspring at *t* being produced by the *t* − 1 individuals that survive to the *t* − 1 → *t* transition, with the density dependent ecology being approximated by the average 1+ abundance of the two timesteps:

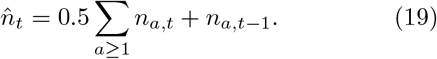

Density regulation is then defining the realised age-structured reproduction and age of maturity

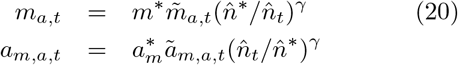

as a log-linear deviation from the intrinsic equilibrium scaled parameters 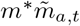 and 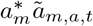, with *γ* Being the strength of regulation, and the number of offspring in age-class zero being

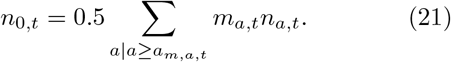

As it is the fitness variation among the quality variants in the population that imposes natural selection in my model, I use the population level model of eqn 13 to turn the intra-population phenotypic variation (denoted by subscript *i*) in quality and fitness into a population level response that determines the average competitive quality of the offspring in age-class zero. For this I follow Witting (1997, 2000b) and define the intra-population variation in relative fitness as

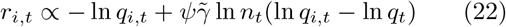

with the variant specific quality term (− ln *q*_*i,t*_) representing the influence of the quality-quantity trade-off (eqn 16) on the growth rate (eqn 13), and the right-hand term of the equation representing the density-frequency-dependent selection of interactive competition, with 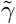 ln *n*_*t*_ being the density dependence in the level of interactive competition, and *ψ* the intra-population differentiation in resource access across the differentiation in competitive quality (ln *q*_*i,t*_ ln *q*_*t*_; *q*_*t*_ is average quality at timestep *t*) per unit interactive competition. This generates the selection gradient

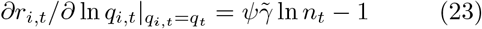

and the following proportional selection response

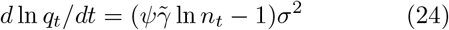

as an approximation for weak and moderate selection following the logic of the secondary theorem of natural selection (Robertson 1968; Taylor 1996), where the additive heritable variance 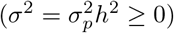 is the phenotypic variance 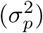 of ln *q* multiplied by the narrow-sense heritability (*h*^2^) of the inclusive inheritance system (Mameli 2004; Danchin et al. 2011), with the additive heritable variance decomposed into a series of additive inheritance terms 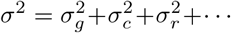 where 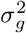 is the additive genetic variance, 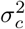 the additive variance of cultural inheritance, 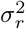 a potential plastic response where the cultural inheritance of offspring is adjusted by an additive proportional plastic response to the selection pressure, and … represents additional unspecified inheritance terms, following the logic of Findlay (1992), Witting (2000b), and Okasha (2007) for multiple additive inheritance components. This provides a proportional selection response that is assumed to apply as an approximation over the limited span of cyclic life history changes necessary for the estimated trajectories.

The response of eqn 24 transforms into a multiplicative formulation

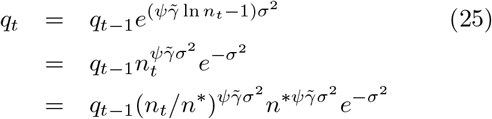

that reduces to

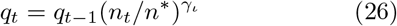

as eqn 23 defines the selection determined equilibrium abundance as 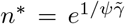, with 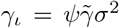 being the selection response parameter that integrates the density dependence of interactive competition 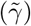 with the resource availability selection (*ψ*) of the density-frequency-dependent interactive competition and the additive heritable variance (*σ*^2^) response of the inclusive inheritance system to the selection. Focussing on the between generation response of selection, I approximate average offspring quality

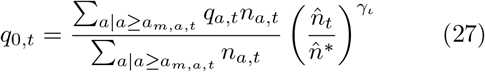

as the average quality of the mature component multiplied by the density-frequency-dependent selection. There is also a potential cohort selection from differential mortality, but I do not consider this selection as I focus on differentiated reproduction imposed by the resource availability of interactive competition.

As I keep the species-specific equilibrium demo-graphic models of Witting (2024a) constant, I am only estimating three parameters (*n*^*^, *γ, γ*_*ι*_) and two initial conditions (*n*_*t*=0_, *q*_*t*=0_) from the abundance data. For the initial conditions of an iteration, I use the same quality across all individuals and distributes the initial abundance according to the stable age-structure

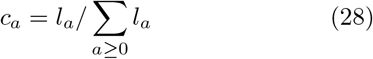

where *l*_0_ = 1, *l*_*a*_ = *p*_0_ *p*^*a−*1^ for 1 ≤ *a < x*, and *l*_*x*_ = *p*_0_ *p*^*x−*1^*/*(1 − *p*).

### A.2 Selection regulated dynamics

Witting (1997, 2000b) describes the population dynamic behaviour of the discrete selection-regulated model with non-overlapping generations (eqn 1 of main paper). This model has damped population cycles when *γ*_*ι*_ *< γ*, neutrally stable cycles when *γ*_*ι*_ = *γ*, and repelling cycles when *γ*_*ι*_ *> γ*. The population period of the stable cycles increases from four to an infinite number of generations as the *γ*_*ι*_ = *γ* parameters decline from two to zero. For a given *γ* the period increases with a decline in *γ*_*ι*_, i.e., with an increasingly damped cycle. When, for a stable cycle, *γ*_*ι*_ = *γ* increases from two to four, there is an extra period in the amplitude of the population period, with the latter declining monotonically to two generations, with the dynamics becoming chaotic when *γ*_*ι*_ = *γ* increases beyond four.

The age-structured model with overlapping generations behaves in a somewhat similar way, but the dynamics depend on the age of reproductive maturity (*a*_*m*_) and the reproductive period [*t*_*r*_ = 1*/*(1 − *p*)]. The age-structured model converges on the discrete model as *a*_*m*_ → 1, *t*_*r*_ → 1, and *p* → 0. With no regulation on maturity, the period (*T*) of the stable population cycle remains a declining function of *γ* (Fig. 3b), with the slope/exponent (*β*) of the ln *T* ∝ *β* ln *γ* relation being −0.5 (estimated by linear regression). The cyclic dynamics become more and more stable with a decline in *γ*_*ι*_, but the damping is also dependent on *a*_*m*_ and *t*_*r*_. The stable cycle, e.g., has a *γ*_*ι*_*/γ* ratio that increases beyond unity as *a*_*m*_ and *t*_*r*_ increase above unity (Fig. 3a and c). For any given combination of *a*_*m*_ and *t*_*r*_, the stable cycle has a *γ*_*ι*_*/γ* ratio that is almost constant (Fig. 3a).

**Figure 3:**
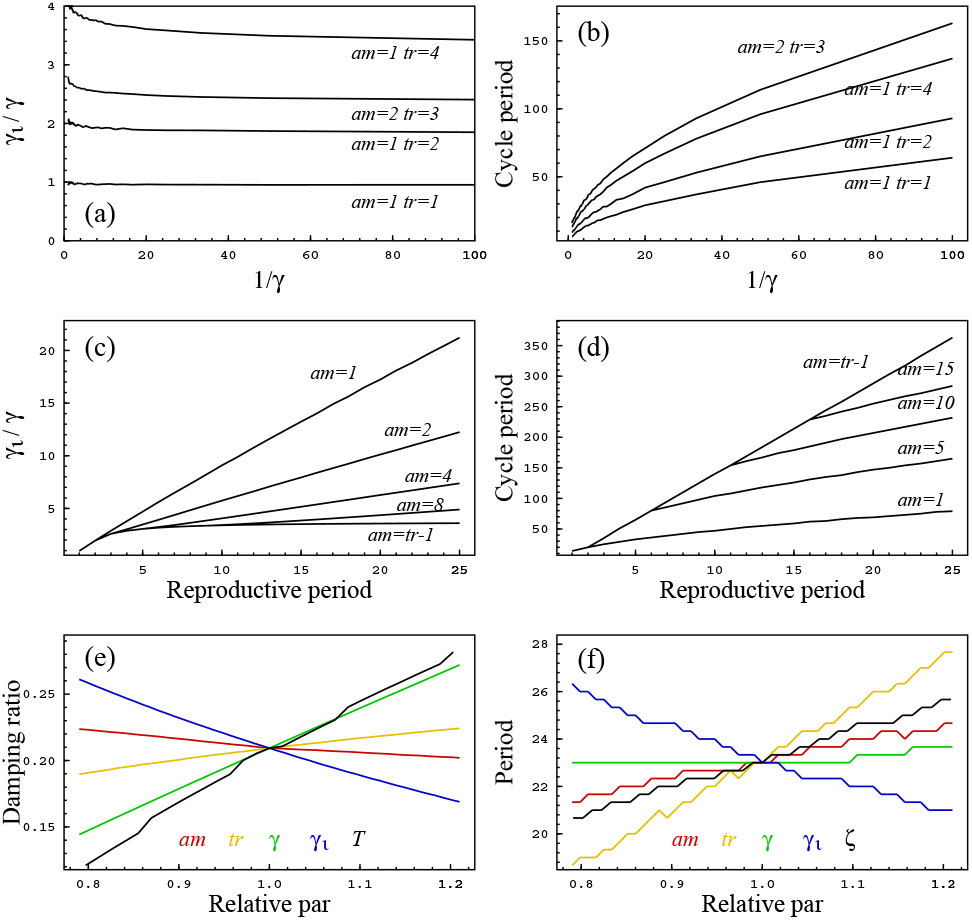
Dynamic behaviour. Plot a to d: The *γ*_*ι*_/*γ*-ratio, and period (in years), of a stable population cycle (ζ = 0) as a function of 1/γ (plot **a** and **b**) and the reproductive period (plot **c** and **d** for γ = 0.2), for different combinations of *tm* and *tr*. **Plot e and f:** The damping ratio (ζ) and population period (*T*) as a function of the parameters *x* ∈ {*am, tr, γ, γ*_*ι*_, ζ, *T*}, relative 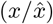 to 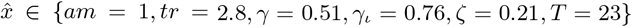. The dependence on *T* in plot **e**, and on ζ in plot **f**, is given by their responses to changes in *γ*_*ι*_.

For a given *γ*, the period of the stable population cycle increases almost linearly with an increase in *a*_*m*_ and *t*_*r*_ (Fig. 3d), with the period dependence on *γ* being somewhat elevated relative to the discrete model where *a*_*m*_ = *t*_*r*_ = 1 (Fig. 3b). Hence, for populations where *γ* is independent of *a*_*m*_ and *t*_*r*_, we can expect an approximate linear relation between the population period *T* and life history periods like *a*_*m*_ and *t*_*r*_.

When only one parameter is altered at the time, the period is almost invariant of *γ* (Fig. 3f). This reflects that the decline in period with an increase in *γ* for dynamics with a given damping ratio, is counterbalanced by the increase in period that is caused by the increased stability of the cycle, as the *γ*_*ι*_*/γ* ratio—that defines the damping ratio—declines with the increase in *γ*. For single parameter perturbations, the damping ratio is usually most strongly dependent on *γ* and *γ*_*ι*_, showing only a small increase with *t*_*r*_ and a small decline with an increase in *a*_*m*_ (Fig. 3e).

To describe the cyclic dynamics, I calculated the cycle period (*T*, in generations) and damping ratio (*ζ*). The damping ratio is zero for a stable cycle, increasing to unity for the monotonic return of density regulated growth. I calculated the damping ratio

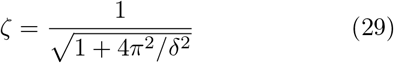

by the logarithmic decrement *δ* = ln(*n*_*p*,1_*/n*_*p*,2_) of the two successive abundance peaks (*n*_*p*,1_ and *n*_*p*,2_) that follow from an equilibrium population that is initiated with a positive growth rate where *q*_*a,t*_ = 2*q*^*^*/*3. The estimated period (*T*) is the number of generations be-ween these two abundance peaks.

When the *γ*_*ι*_*/γ*-ratio increases above one the dynamics become unstable with amplitudes that increase over time instead of dampening out. In these cases, I revert *n*_*p*,1_ and *n*_*p*,2_ in the estimate of *δ* = ln(*n*_*p*,2_*/n*_*p*,1_) and multiply the ratio by minus one, so that negative *ζ* values refer to exploding cycles, with the rate of explosion increasing as *ζ* declines from zero to minus one.

### A.3 Model fitting & model selection

I use the likelihood (*L*) maximum to estimate the parameters of all models given log normally distributed abundance data

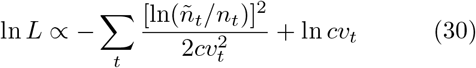

where *ñ*_*t*_ is the 1+ index estimate in year *t, n*_*t*_ is the corresponding model estimate, and the coefficient of variation of the index estimate on normal scale 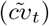 used in an approximation of the *σ* parameter of the log normal distribution [as *σ*^2^ = ln(1 + *cv*^2^)], with 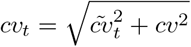 including an additional variance term (*cv*^2^) of the data relative to the model projection, with *cv* estimated by the likelihood function (an additional ln *ñ*_*t*_ term is not included explicitly in the likelihood as it is redundant).

I project each model for 100,000 random parameter sets and apply a Quasi-Newtonian minimiser to find a local likelihood maximum for the 100 best fits, with the global maximum being the maximum across the local maxima. I convert the global likelihood maximum of each model to AIC [*α* = 2(*k* − ln *L*), *k* nr. of model parameters] to select the best model for each timeseries, calculating the fraction of the AIC selected models that include selection.

I use models with fitted additional variance (*cv*) in AIC selection. Relative to a case with no additional variance, this increases the relative likelihood of models with a poor fit, increasing their AIC-selection likelihood. Given that the non-selection models most often have the poorest fit to the abundance data, the exponential and density regulated models are slightly over-represented among the AIC-selected models relative to a case with no additional variance, and this makes my conclusion on the importance of including selection into population dynamic modelling somewhat conservative.

As I focus on the non-fluctuating population dynamics of weak to moderate regulation, I place an upper limit of 1.5 on the estimates of *γ* and *γ*_*ι*_ to avoid the estimation of fluctuating dynamics from between-year zig-zag variation in abundance estimates. To reduce other confounding effects, I use a minimum fitting criterion and analyse fitted models further only if the mean of the residuals is not significantly different from zero (*p <* 0.05 student’s t), there is no significant autocorrelation in the residuals (lag 1 and 2), no significant correlation between the residuals and the model, and the model explains at least 50% of the variance.

A second model-selection includes five selection-regulated models. In addition to the original model with a stable equilibrium abundance, it includes four versions with a linear trend in the population dynamic equilibrium (*n*^*^): *i*) a trend that covers the whole data period [1 extra parameter (final *n*^*^) fitted by the likelihood]; ii) a trend that starts after the first data year [2 extra parameters: trend starting year and final *n*^*^]; iii) a trend that ends before the last data year [2 extra parameters: final *n*^*^ and trend ending year]; and iv) a trend that starts after the first year and ends before the last year [3 extra parameters: start and end year of trend plus final *n*^*^], with a minimum allowed trend period of five years. Following eye inspection of all fits, I allow AIC to select one year of catastrophic survival for ten populations with an obvious crash in abundance, adding two extra likelihood fitted parameters (year and year-specific 1+ survival). For seven large whale populations I subtract annual catches from the dynamics following Witting (2013; data from https://iwc.int).

## References

Agashe D. (2009). The stabilizing effect of intraspecific genetic variation on population dynamics in novel and ancestral habitats. Am. Nat. 174:255–267.

Akaike H. (1973). Information theory as an extension of the maximum likelihood principle. In: Petrov B. N. Csaki F. (eds). Second International Symposium on Information Theory: Akademiai Kiado, pp 267–281.

Andreassen H. P., Sundell J., Ecke F., Halle S., Haapakoski M., Henttonen H., Huitu O., Jacob J., Johnsen K., Koskela E., Luque-Larena J. J., Lecomte N., Leirs H., Marien J., Neby M., Ratti O., Sievert T., Singleton G. R., Cann J. v., Broecke B. V., Ylonen H. (2021). Population cycles and outbreaks of small rodents: ten essential questions we still need to solve. Oecologia 195:601–622.

Bell G. (2017). Evolutionary rescue. Ann. Rev. Ecol. Evol. Syst. 48:605–627.

Bell G. Gonzalez A. (2009). Evolutionary rescue can prevent extinction following enviromental change. Ecol. Lett. 12:942–948.

Berryman A. A. (1996). What causes population cycles of forest lepidoptera? Trends Ecol. Evol. 11:28–32.

Boonstra R. Hochachka W. M. (1997). Maternal effects and additive genetic inheritance in the collared lemming *Dicrostonyx groenlandicus*. Evol. Ecol. 11:169–182.

Boonstra R. Krebs C. J. (1979). Viability of large- and small-sized adults in fluctuating vole populations. Ecology 60:567–573.

Bossdorf O., Richards C. L., Pigliucci M. (2008). Epigenetics for ecologists. Ecol. Lett. 11:106–115.

Brunner F. S., Deere J. A., Egas M., Eizaguirre C., Raeymaekers J. A. M. (2019). The diversity of eco-evolutionary dynamics: Comparing the feedbacks between ecology and evolution across scales. Funct. Ecol. 33:7–12.

BTO (2022). British Trust for Ornithology. Population trend graphs. https://www.bto.org.

Caswell H. (1989). Life-history strategies. In: Cherrett J. M. (ed). Ecological concepts. The contribution of ecology to an understanding of the natural world: Blackwell Scientific Publications, Oxford, pp 285–307.

Charlesworth B. (1994). Evolution in age-structured populations. 2nd edn. Cambridge University Press, Cambridge.

Chitty D. (1960). Population processes in the voles and their relevance to general theory. Can. J. Zool. 38:99–113.

Chitty D. (1996). Do lemmings commit suicide? Beautiful hypotheses and ugly facts. Oxford University Press, New York.

Christiansen F. B. (1991). On conditions for evolutionary stability for a continuously varying character. Am. Nat. 138:37–50.

Coulson T., Macnulty D. R., Stahler D. R., Vonholdt B., Wayne R. K., Smith D. W. (2011). Modeling Effects of Environmental Change on Wolf Population Dynamics, Trait Evolution, and Life History. Science 334:1275–1278.

Danchin E., Charmantier A., Champagne F. A., Mesoudi A., Pujol B., Blanchet S. (2011). Beyond DNA: integrating inclusive inheritance into an extended theory of evolution. Nature Rev., Genet. 12:475–486.

DOF (2022). Dansk Ornitologisk Forening. Punkttællinger. https://www.dof.dk.

Elton C. S. (1924). Periodic fluctuations in number of animals: their causes and effects. Brit. J. Exp. Biolo. 2:119–163.

Eshel I. (1983). Evolutionary and continuous stability. J. theor. Biol. 103:99–111.

Findlay C. S. (1992). Secondary theorem of natural selection in biocultural populations. Theor. Pop. Biol. 41:72–89.

Fisher R. A. (1930). The genetical theory of natural selection. Clarendon, Oxford.

Ford E. B. (1931). Mendelism and evolution. Methuen, London.

Ginzburg L. R. (1998). Inertial growth. Population dynamics based on maternal effects. In: Mousseau T. A. Fox C. W. (eds). Maternal effects as adaptations: Oxford University Press, New York, pp 42–53.

Gomulkiewicz R. Holt R. D. (1995). When does evolution by natural selection prevent extinction? Evolution 49:201–207.

Haigh J. Rose M. R. (1980). Evolutionary game auctions. J. theor. Biol. 85:381–397.

Hairston J. N. G. Hairston S. N. G. (1993). Cause-effect relationships in energy flow, trophic structure, and interspecific interactions. Am. Nat. 142:379–411.

Hairston N. G. J., Ellner S. P., Geber M. A., Yoshida T., Fox J. A. (2005). Rapid evolution and the convergence of ecological and evolutionary time. Ecol. Lett. 8:1114–1127.

Hairston S. N. G., Smith F. E., Slobodkin L. B. (1960). Community structure, population control, and competition. Am. Nat. 94:421–425.

Halley J. M., Vokou D., Pappas G., Sainis I. (2021). SARS-CoV-2 mutational cascades and the risk of hyper-exponential growth. Microbial Pathigenesis 161:10.1016/j.micpath.2021.105237.

Hansen T. F., Stenseth N. C., Henttonen H., Tast J. (1999). Interspecific and intraspecific competition as causes of direct and delayed density dependence in a fluctuating vole population. Proc. Nat. Acad. Sci. USA 96:986–991.

Hardy I. C. W. Briffa M. (2013). Animal contests. Cambridge University Press, Cambridge.

Hendry A. P. (2017). Eco-evolutionary dynamics. Princeton University Press, Princeton.

Hodges K. E., Stefan C. I., Gillis E. A. (1999). Does body condition affect fecundity in a cyclic population of snowshoe hares? Can. J. Zool. 77:1–6.

Holt S. J. (2004). Counting whales in the North Atlantic. Science 303:39.

Hörnfeldt B. (1994). Delayed density dependence as a determinant of vole cycles. Ecology 73:791–806.

Inchausti P. Ginzburg L. R. (2009). Maternal effects mechanism of population cycling: a formidable competitor to the traditional predator. Phil. Trans. R. Soc. B: Biol. Sci 364:1117–1124.

Kaitala V., Ranta E., Lindstrom J. (1996). Cyclic population dynamics and random perturbations. J. Anim. Ecol. 65:249–251.

Keith L. B. (1963). Wildlife’s ten year cycle. University of Wisconsin Press, Madison.

Knape J. deValpine P. (2012). Are patterns of density dependence in the Global Population Dynamic Database driven by uncertainty about population abundance? Ecol. Lett. 15:17–23.

Knaus P., Schmid H., Strebel N., Sattler T. (2022). The State of Birds in Switzerland 2022 online. http://www.vogelwarte.ch.

Koenig W. D. (2002). Global patterns of environmental synchrony and the Moran effect. Ecography 25:283–288.

Le Boeuf B. J. Briggs K. T. (1977). The cost of living in a seal harem. Mammalia 41:167–195.

Lidicker W. Z. Ostfeld R. S. (1991). Extra-large body size in California voles: Causes and fitness consequences. Oikos 61:108–121.

Liebhold A., Koenig W. D., Bjørnstad O. N. (2004). Spatial synchrony in population dynamics. Ann. Rev. Ecol. Evol. Syst. 35:467–490.

Liu R., Gourley S. A., DeAngelis D. L., Bryant J. P. (2013). A mathematical model of woody plant chemical defenses and snowshoe hare feeding behavior in boreal forests: the effect of age-dependent toxicity of twig segments. SIAM J. Appl. Math. 73:281–304.

LPI (2022). Living Planet Index database. www.livingplanetindex.org.

Malthus T. R. (1798). An essay on the principle of population. Johnson, London.

Mameli M. (2004). Nongenetic selection and nongenetic inheritance. Brit. J. Phil. Sci. 55:35–71.

May R. M. (2020). What determines population density?. In: Dobson A., Holt R. D., Tilman D. (eds). Unsolved problems in ecology: Princeton University Press, Princeton, pp 67–75.

Maynard Smith J. (1964). Group selection and kin selection. Nature 201:1145–1146.

Maynard Smith J. (1982). Evolution and the theory of games. Cambridge University Press, Cambridge.

McCauley E., Nelson W., Nisbet R. (2008). Small-amplitude cycles emerge from stage-structured interactions in daphnia algal systems. Nature 455:1240–1243.

McCullough D. R. (1979). The George River Deer Herd: Population ecology of a k-selected species. Univ. Michigan Press, Ann Arbor.

McKane A. J. Newman T. J. (2005). Predator-prey cycles from resonant amplification of demographic stochasticity. Phys. Rev. Lett. 94:218102.

Murdoch W. W., Kendall B. E., Nisbet R. M., Briggs C. J., Mccauley E., Bolser R. (2002). Single-species models for many-species food webs. Nature 417:541–543.

Murdoch W. W. McCauley E. (1985). Three distinct types of dynamic behavior shown by a single planktonic system. Nature 316:628–630.

Myers J. H. (1990). Population cycles of western tent caterpillars: experimental introductions and synchrony of fluctuations. Ecology 71:986–995.

Myers J. H. (2018). Population cycles: generalities, exceptions and remaining mysteries. Proc. R. Soc. B. 285:20172841.

Naumov S. P., Gibet L. A., Shatalova S. (1969). Dynamics of sex ratio in respect to changes in numbers of mammals. Zh. Obshch. Biol. 30:673–680.

Norrdahl K. Korpimäki E. (2002). Changes in individual quality during a 3-year population cycle of voles. Oecologia 130:239–249.

Okasha S. (2007). Cultural inheritance and Fisher’s “fundamental theorem” of natural selection. Biol. Theory 2:290–299.

Oli M. K. (2019). Population cycles in voles and lemmings: state of the science and future directions. Mamm. Rev. 49:226–239.

Parker G. A. (1983). Arms races in evolution–an ess to the opponent-independent cost game. J. theor. Biol. 101:619–648.

Pavithran I. Sujith R. I. (2022). Extreme COVID-19 waves reveal hyperexponential growth and finite-time singularity. Chaos 32:041104.

PECBMS (2022). Pan-European Common Bird Monitoring Scheme. https://pecbms.info.

Pella J. Tomlinson P. (1969). A generalized stock production model. Trop. Tuna. Comm. Bull. 13:419–496.

Pfennig, D. W., ed (2021). Phenotypic plasticity & evolution: causes, consequences, controversies. CRC Pres, Oxon.

Piertney S. B., Lambin X., Maccoll A. D. C., Lock K., Bacon P. J., Dallas J. F., Leckie F., Mougeot F., Racey P. A., Redpath S., Moss R. (2008). Temporal changes in kin structure through a population cycle in a territorial bird, the red grouse *Lagopus lagopus scoticus*. Mol. Ecol. 17:2544–2551.

Post E. Forchhammer M. C. (2002). Synchronization of animal population dynamics by large-scale climate. Nature 420:168–171.

Ramsayer J., Kaltz O., Hochberg M. E. (2013). Evolutionary rescue in populations of *Peudomonas fluoresens* across an antibiotic gradient. Evol. Appl. 6:608–616.

Ranta E., Kaitala V., Lindstrom J., Linden H. (1995). Synchrony in population dynamics. Proc. R. Soc. Lond. B. 262:113–118.

Richards E. J. (2006). Inherited epigenetic variation – revisiting soft inheritance. Nature Rev., Genet. 7:395–401.

Robertson A. (1968). The spectrum of genetic variation. In: Lewontin R. C. (ed). Population Biology and Evolution: Syracuse University Press, New York, pp 5–16.

Roff D. A. (1992). The evolution of life histories. Theory and analysis. University of Chicago Press, New York.

Roughgarden J. (1971). Density-dependent natural selection. Ecology 5:453–468.

Saccheri I. Hanski I. (2006). Natural selection and population dynamics. Trends Ecol. Evol. 21:341–347.

Sauer J. R., Niven D. K., Hines J. E., Ziolkowski D. J., Pardieck K. L., Fallon J. E., Link W. A. (2017). The North American Breeding Bird Survey, Results and analysis 1996 – 2015. Version 2.07.2017. USGS Patuxent Wildlife Research Center, Laurel, Maryland, Available at www.mbr-pwrc.usgs.gov/bbs/bbs.html.

SFT (2022). Svensk Fågeltaxering. http://www.fageltaxering.lu.se.

Sibly R. M., Baker D., Denham M. C., Hone J., Pagel M. (2005). On the regulation of populations of mammals, birds, fish, and insects. Science 309:607–610.

Simchuk A. P., Ivashov A. V., Companiytsev V. A. (1999). Genetic patters as possible factors causing population cycles in oak leafroller moth, *Tortrix viridana* L. For. Ecol. Manage. 113:35–49.

Simpson G. G. (1953). The major features of evolution. Columbia University Press, New York.

Sinclair A. R. E. (1989). Population regulation in animals. In: Cherrett J. M. (ed). Ecological concepts. The contribution of ecology to an understanding of the natural world: Blackwell Scientific Publications, Oxford, pp 197–241.

Sinervo B., Svensson E., Comendant T. (2000). Density cycles and an offspring quantity and quality game driven by natural selection. Nature 406:985–988.

Smith C. C. Fretwell S. D. (1974). The optimal balance between size and number of offspring. Am. Nat. 108:499–506.

Snell-Rood E. C., Kobiela M. E., Sikkink K. L., Shephard A. M. (2018). Mechanisms of plastic rescue in novel environments. Ann. Rev. Ecol. Evol. Syst. 49:331–354.

Sovon (2022). Netwerk Ecologische Monitoring, Sovon. Provincies & CBS. http://sovon.nl.

Stearns S. C. (1992). The evolution of life histories. Oxford University Press, Oxford.

Stenseth N. C. (1981). On chitty’s theory for fluctuating population: the importance of genetic polymorphism in the generation of regular density cycles. J. theor. Biol. 90:9–36.

Stenseth N. C. (1985). Mathematical models of microtine cycles: models and the real world. Acta Zool. Fenn. 173:7–12.

Taylor P. D. (1989). Evolutionary stability in one-parameter models under weak selection. Theor. Pop. Biol. 36:125–143.

Taylor P. D. (1996). The selection differential in quantitative genetics and ess models. Evolution 50:2106–2110.

Taylor R. A., White A., Sherratt J. (2013). How do variations in seasonality affect population cycles? Proc. R. Soc. Lond. B. 10.1098/rspb.2012.2714.

Thompson J. N. (1998). Rapid evolution as an ecological process. Trends Ecol. Evol. 13:329–332.

Turchin P. Taylor A. D. (1992). Complex dynamics in ecological time series. Ecology 73:289–305.

Turcotte M. M., Reznick D. N., Hare J. D. (2011). The impact of rapid evolution on population dynamics in the wild: experimental test of eco-evolutionary dynamics. Ecol. Lett. 14:1084–1092.

Tyson R., Haines S., Hodges K. (2010). Modelling the Canada lynx and snowshoe hare population cycle: the role of specialist predators. Theor. Ecol. 3:97–111.

Vermeij G. J. (1987). Evolution and escalation. Princeton University Press, Princeton.

Voipio P. (1950). Evolution at the population level with special reference to game animals and practical game management. Papers Game Res. 5:1–176.

Voipio P. (1988). Comments on the implications of genetic ingredients in animal population dynamics. Acta Zool. Fenn. 25:321–333.

Whitehead H., Laland K. N., Rendell L., Thorogood R., Whiten A. (2019). The reach of gene-culture coevolution in animals. Nature Comm. 10:2405.

Wiens J. A. (1966). On group selection and wynne-edwards’ hypothesis. Am. Sci. 54:273–287.

Williams G. C. (1966). Adaptation and natural selection. A critique of some current evolutionary thought. Princeton University Press, Princeton.

Witting L. (1995). The body mass allometries as evolutionarily determined by the foraging of mobile organisms. J. theor. Biol. 177:129–137, 10.1006/jtbi.1995.0231.

Witting L. (1997). A general theory of evolution. By means of selection by density dependent competitive interactions. Peregrine Publisher, Århus, 330 pp, URL https://mrLife.org.

Witting L. (2000a). Interference competition set limits to the fundamental theorem of natural selection. Acta Biotheor. 48:107–120, 10.1023/A:1002788313345.

Witting L. (2000b). Population cycles caused by selection by density dependent competitive interactions. Bull. Math. Biol. 62:1109–1136, 10.1006/bulm.2000.0200.

Witting L. (2002a). Evolutionary dynamics of exploited populations selected by density dependent competitive interactions. Ecol. Model. 157:51–68, 10.1016/S0304–3800(02)00172–2.

Witting L. (2002b). From asexual to eusocial reproduction by multilevel selection by density dependent competitive interactions. Theor. Pop. Biol. 61:171–195, 10.1006/tpbi.2001.1561.

Witting L. (2008). Inevitable evolution: back to *The Origin* and beyond the 20th Century paradigm of contingent evolution by historical natural selection. Biol. Rev. 83:259–294, 10.1111/j.1469–185X.2008.00043.x.

Witting L. (2013). Selection-delayed population dynamics in baleen whales and beyond. Pop. Ecol. 55:377–401, 10.1007/s10144–013–0370–9.

Witting L. (2017a). The natural selection of metabolism and mass selects allometric transitions from prokaryotes to mammals. Theor. Pop. Biol. 117:23–42, 10.1016/j.tpb.2017.08.005.

Witting L. (2017b). The natural selection of metabolism and mass selects lifeforms from viruses to multicellular animals. Ecol. Evol. 7:9098–9118, 10.1002/ece3.3432.

Witting L. (2023a). On the natural selection of body mass allometries. Acta Oecol. 118:103889, 10.1016/j.actao.2023.103889.

Witting L. (2023b). Population dynamic population delineation in North American birds. Preprint at bioRxiv 10.1101/2023.08.29.555290.

Witting L. (2024). Population dynamic life history models of the birds and mammals of the world. Ecol. Info. 80:102492.

Witting L. (2026). Self-organising natural selection from replicating molecules to multicellular sexually reproducing organisms. Preprint at EcoEvoRxiv 10.32942/X26G8B.

Wolda H. Dennis B. (1993). Density-dependence tests, are they. Oecologia 95:581–591.

Wynne-Edwards V. (1959). The control of population density through social behaviour: A hypothesis. Ibis 101:436–441.

Wynne-Edwards V. C. (1993). A rationale for group selection. J. theor. Biol. 162:1–22.

Yoshida T., Jones L. E., Ellner S. P., Fussmann G. F., Hairston N. G. (2003). Rapid evolution drives ecological dynamics in a predator-prey system. Nature 424:303–306.

